# Hydroxide ions amplified by water entanglement underly the mechanism of general anesthesia

**DOI:** 10.1101/2021.01.28.428716

**Authors:** Hao Qian, Na Li, Lei Yang, Younian Xu, Rong Chen, Dongshi Lu, Ruihan Zhao, Hui Liu, Nanxue Cui, Qiao Zhou, Shihai Zhang

**Author notes:** These authors contributed equally to this work (Hao Qian, Na Li, Lei Yang, Younian Xu).

## Abstract

It is believed that inhaled anesthetics occupy hydrophobic pockets within target proteins, but how inhaled anesthetics with diverse shapes and sizes fit into highly structurally selective pockets is unknown. For hydroxide ions are hydrophobic, we determined whether hydroxide ions could bridge inhaled anesthetics and protein pockets. We found that small additional load of cerebral hydroxide ions decreases anesthetic potency. Multiple-water entanglement network, derived from Ising model, has a great ability to amplify ultralow changes in the cerebral hydroxide ion concentration, and consequently, amplified hydroxide ions account for neural excitability. Molecular dynamics simulations showed that inhaled anesthetics produce anesthesia by attenuating the formation of multiple-water entanglement network. This work suggests amplified hydroxide ions underlying a unified mechanism for the anesthetic action of inhaled anesthetics.

---

General anesthesia is indispensable. Millions of surgical procedures and invasive examinations are performed under general anesthesia each year. General anesthesia is also fascinating because general anesthesia has been alluded to be an experimental inroad to consciousness (*1-4*) which is perhaps the biggest mystery in nature, based on the facts that general anesthesia, a reversibly drug-induced unconscious state, is definable and anesthetic potency is quantifiable. Although general anesthesia is both indispensable and fascinating, the mechanism of action of inhaled anesthetics (IAs) remains obscure over 170 years (*5, 6*). The challenge, largely in part, is what property a host of molecules with highly diverse chemical structures can possibly have in common that causes general anesthesia (*2*). Since at the fundamental level all things are quantum, quantum processing may remove the diversities in chemical structure specific to individual anesthetics. Quantum properties may thus provide a corollary answer to unify diverse IAs into a common quantum-processing mechanism of anesthetic action (*3, 7*). Indeed, the notion that the element hosting consciousness must have a nuclear spin of ½ has been proposed (*8*), based on the fact that singlets formed by *I* = nuclei are long-lasting because they are immune to the electric fields that neurons produce in the brain. This proposal has been confirmed, albeit indirectly, by our previous experiment (*3*) on the isotopic dependence of xenon anesthesia: ^129^xenon (*I* = ½) is less potent than xenon isotopes with zero nuclear spin, strongly indicating that the nuclear-spin property of ^129^xenon partly antagonizes its anesthetic property, suggesting a quantum-processing effect.

How is the nuclear spin involved in the anesthetic action of xenon anesthesia, and could quantum processing be extended from xenon to all IAs? If there exists a component in the brain that accounts for neural excitability through quantum processing with nuclear spins, these questions are answered, as IAs inhibit neural excitability by reducing this component explicitly as general anesthesia. As such, the candidate component must have three properties—crucial importance in all living organisms (*9*), nuclear spin or total nuclear spin of ½, and a size small enough to exert quantum effects. Phosphorus and hydrogen are the only two biologically prevalent light elements with *I* = ½. Phosphorus has been experimentally ruled out (*10*), leaving hydrogen as the only candidate. In living systems, most hydrogens are within water molecules. The importance of liquid water for the existence of life on Earth is undeniable. The water dissociation reaction, 2H_2_O → H_3_O^+^ + OH^-^, generates a pair of hydronium (H_3_O^+^) and hydroxide (OH^-^) ions, both having total nuclear spins of ½, both small enough to exert quantum effects, and both populating biochemistry. Using the 50% effective dose for loss of righting reflex (LORR ED_50_) for mice to sevoflurane as a measure, we report here that cerebral OH^-^ ions amplified by a multiple-water entanglement network (MWEN) are necessary for neural excitability, and attenuating amplification is the mechanism of action of IAs.

## LORR ED_50_ measurement

To gain insight into how H_3_O^+^ and OH^-^ ions may affect anesthetic potency we examined the effects of additional loads of H_3_O^+^ and OH^-^ ions by injecting weak acids or bases into mouse lateral ventricle on LORR ED_50_ for C57BL/6 male mice (aged 8 weeks) to a typical potent IA, sevoflurane. For comparison of the effects of equal amounts of overloaded H_3_O^+^ or OH^-^ ions on the anesthetic potency of sevoflurane, the pH values of the weak acids ammonium chloride and heparin sodium (*11*), were both 6.25, and those of the weak bases sodium acetate and xanthohumol (*11*), were both 7.75, with injection volumes of 2 μl. Our method was effective to determine drug-induced anesthetic effects (fig. S1), for the mice undergoing intracerebroventricular cannulation were regarded as intact animals (*10*). We found that injections of two controls, 0.9% NaCl and ammonium acetate, did not affect the LORR ED50 (Fig. 1, A and B). Ammonium chloride (Fig. 1C) and heparin sodium (Fig. 1D) decreased the ED_50_. However, sodium acetate (Fig. 1E) and xanthohumol (Fig. 1F) increased the ED_50_. Because an increased ED_50_ means decreased potency of an IA and *vice versa* (*12*), these findings indicate that, regardless of the chemical structure, weak acids enhance whereas weak bases weaken the anesthetic potency of sevoflurane by changing the cerebral pH in opposite directions.

**Fig. 1.**
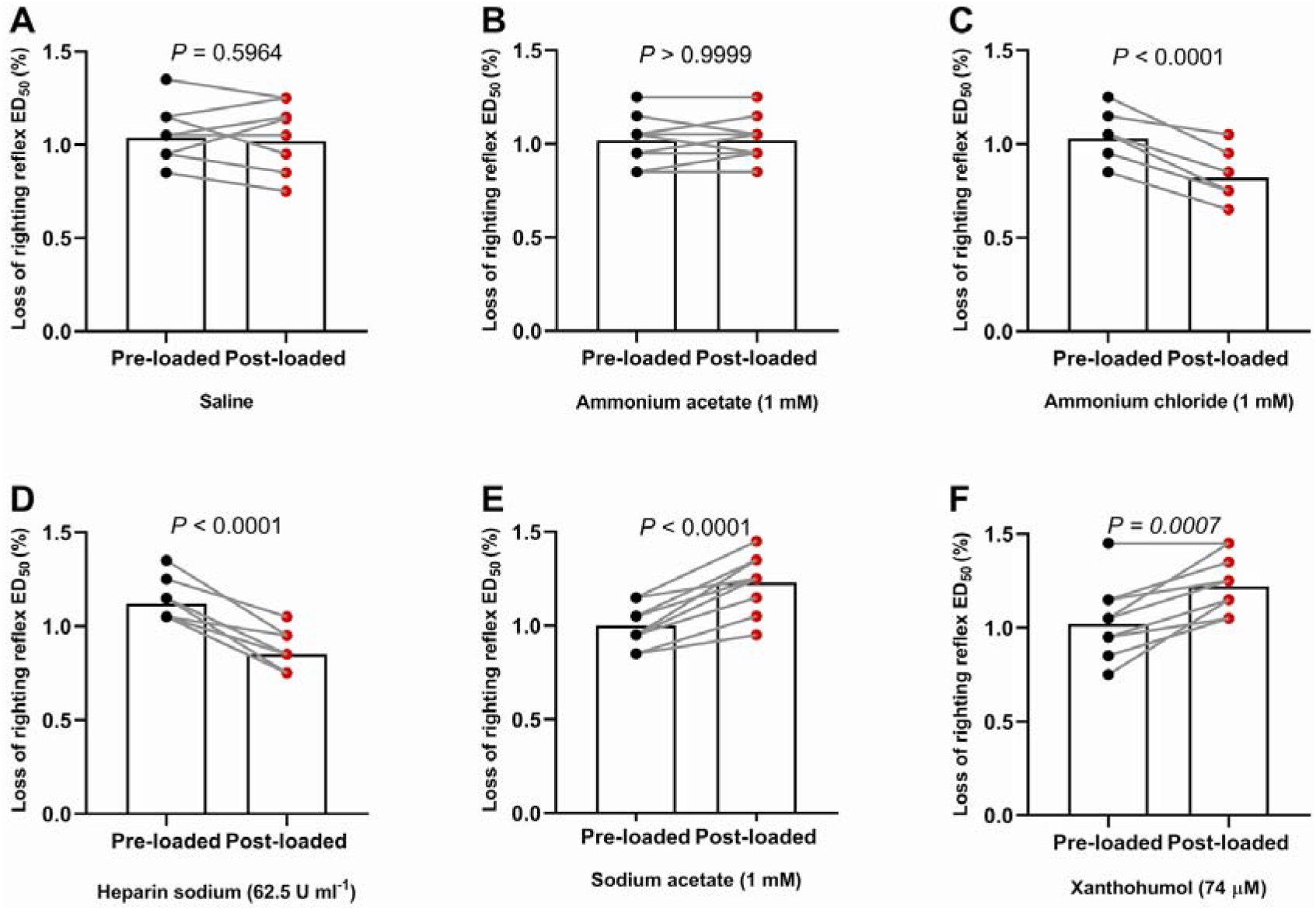
Effects of weak acids and weak bases on the LORR ED_50_ of sevoflurane in mice. Saline and weak acids and bases were intracerebroventricularly injected into mice. **A**, 0.9% NaCl did not affect the LORR ED_50_ (pre-SD, 0.15, post-SD, 0.19; and 95% CI, 0.93 to 1.15% versus 0.88 to 1.15%). **B**, Ammonium acetate had no effect (pre-SD, 0.13, post-SD, 0.13; and 95% CI, 0.93 to 1.11% versus 0.94 to 1.10%). Weak acids reduced the LORR ED_50_ values: **C**, ammonium chloride (pre-SD, 0.11, post-SD, 0.12; and 95% CI, 0.95 to 1.11% versus 0.74 to 0.90%) and **D**, heparin sodium (pre-SD, 0.11, post-SD, 0.11; and 95% CI, 1.04 to 1.20% versus 0.77 to 0.92%). Weak bases increased the LORR ED_50_ values: **E**, sodium acetate (pre-SD, 0.15, post-SD, 0.13; and 95% CI, 1.11 to 1.32% versus 0.92 to 1.10%) and **F**, xanthohumol (pre-SD, 0.18, post-SD, 0.16; and 95% CI, 0.88 to 1.16% versus 1.11 to 1.33%). The injection volume was 2 μl. Mean (*n* = 10 mice), two-tailed paired *t*-test. The LORR ED_50_ is presented as the sevoflurane concentration (%). LORR ED_50_ denotes the 50% effective dose for the loss of righting reflex.

However, the net changes in pH in the mouse brain (*13*) resulting from the intracerebroventricular injections of weak acids and bases were as small as 0.001 pH units or even less, which were equivalent to a net increase or decrease of 137 free H_3_O^+^ or OH^-^ ions in a pyramidal cell, for example (supplementary text). How could the ultralow changes in cerebral H_3_O^+^ or OH^-^ ions affect anesthetic potency so considerably? Apparently, there should exist a system which can amplify the ultralow changes in cerebral H_3_O^+^ or OH^-^ ions for a considerable manifestation of anesthetic effects. Quantum mechanics has an innate characteristic of amplification (*14, 15*). In living system, quantum amplification has been found to function in avian compasses (*16*).

### MWEN-mediated amplification

The existence of MWEN has been repeatedly implied (*17-20*) and been observed in liquid water (*21*). We postulated that an MWEN may be the system able to amplify H_3_O^+^ and OH^-^ ions.

To verify the postulation, we first formulated a scheme for an MWEN using one-dimensional Ising model, based on the fact that both H_3_O^+^ and OH^-^ ions have total nuclear spins of ½. In liquid water, electric field fluctuations result in H_2_O molecules dissociating and subsequently drive nascent H_3_O^+^ and OH^-^ ions to move rapidly away from each other to a distance too far for them to reinstate into their initial H_2_O molecule because the electrostatic attraction between them fades (*22*). Therefore, it is most likely for the pair of ions to combine with nearby counterions dissociated from other H_2_O molecules. Because the two hydrogen atoms in a H_2_O molecule (ortho-or para-H_2_O) are indistinguishable and nuclear-spin entangled, the nascent H_3_O^+^ and OH^-^ ions inherit the indistinguishability and entanglement from their initial H_2_O molecule, analogous to the nuclear-spin entanglement between the two separated phosphorus ions cleaved from a diphosphate molecule (*8*).

According to the method described by Teng and Sy (*23*), we derived a 1-d Ising model to describe that the spins of H_3_O^+^ and OH^-^ ions are naturally entangled.

Because the total nuclear spins of H_3_O^+^ and OH^-^ ions are of ½, the Hamiltonian of the Ising chain is

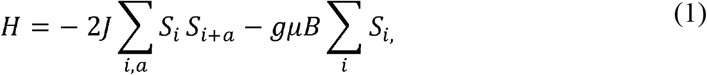

where *J, g* and *μ* have the traditional meanings, *S_i_* is the spin for the *i*th ion in the spin lattice, *i* and *i+a* are indices labelling the adjacent ions, and *B* is the magnitude of the external magnetic field that can align the spins along its direction irrespective of the temperature (*24*).

According to Green’s function, the function of the site-spin (with *B* ≠ 0) for the Ising chain (*24*) is

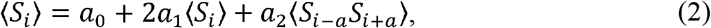

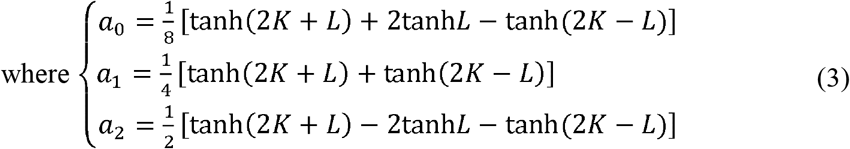

and *L* = *gμB/2kBT* and *K* = *J/2kBT*.

In the absence of external magnetic field, for the two-site spin correlations we have

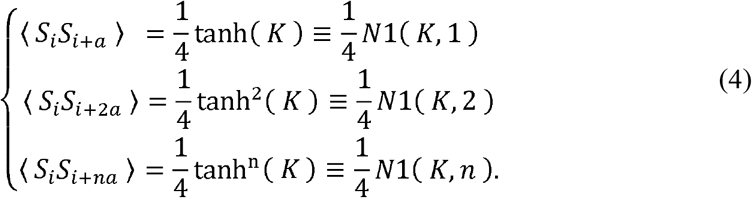

Evidently, there always exists

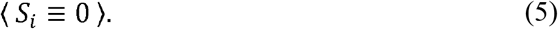

Apparently, equation 4 requires that, in order to keep the average values of the two-site spins to be ¼, the spin configuration must be in alignment. Equation 5 requires that, in order to maintain the average value of the single-site spin to be zero, the number of up-spins must equal that of down-spins. Particles are entangled in the Ising chain if and only if they fulfil the two requirements. Do H_3_O^+^ and OH^-^ ions fulfill the requirements? The nuclear spin configuration of hydrogen nuclei within ortho-or para-water molecules is in alignment, which is inherited by H_3_O^+^ and OH^-^ ions when water dissociation takes place. Therefore, the spin configuration of H_3_O^+^ and OH^-^ ions fulfils the first requirement. Water dissociations yield equal numbers of H_3_O^+^ and OH^-^ ions. In average, the numbers of up-spins and down-spins are equal. H_3_O^+^ and OH^-^ ions thus fulfill the second requirement as well. Therefore, the nuclear spins of H_3_O^+^ and OH^-^ ions must be entangled in the Ising chain at any finite temperature irrespective of external magnetic field (*23*). The entangled network of H_3_O^+^ and OH^-^ ions is termed as an MWEN for simplification of description.

We next calculated the amplification capacity of an MWEN. In neutral liquid water, H_3_O^+^ and OH^-^ ions are far away from each other, with a dissociation constant of water, *K_w_*, of 1 × 10^-14^ at 25 □. The recombination of H_3_O^+^ and OH^-^ ions under this condition can be effectively described by the Eigen and de Maeyer (*25*) model. According to the model, the lifetime of H_3_O^+^ or OH^-^ ions is 10^-4^ s. To maintain the equilibrium of H_3_O^+^ and OH^-^ ion concentrations in water, neutralization of a pair of H_3_O^+^ and OH^-^ ions produces a new pair at the same time. That is, two numbers are possibly added to the MWEN in every 10^-4^ s interval. We next consider the decoherence of an MWEN. An MWEN is composed of hydrogen nuclei with nuclear spins of ½. The nuclei thus do not couple to electric fields; they interact with the environment only through magnetic fields, *e.g.*, dipole fields from nuclear spins external to the MWEN. Therefore, the primary mechanism for decoherence of the network is entanglement of the hydrogen nuclei spins within the network with external nuclear spins. The decoherence time for the hydrogen spins in an MWEN is the nuclear magnetic resonance (NMR) spin-lattice relaxation time *T*_1_, analogous to the decoherence time of spins of phosphorus nuclei within a Posner molecule (*26*). Given *N* as the number in the MWEN, and *T*_1_ as the decoherence time of any number in the network, the decoherence time, *t_de_*, of the entire network is *t_de_* = *T*_1_*N*. Given *t_H_* the mean lifetime of H_3_O^+^ or OH^-^ ions and *t*_ent_, the time needed for the formation of the entire network, and because two numbers are added to the network every *t*_H_, we have 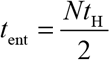. Because *t*_ent_ = *t*_de_, we have 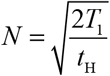 and 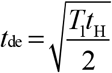. Inserting *T*_1_ =3 s (*27*) and *t*_H_ =10^-4^ s (*25*) into the equations, we obtain roughly *N* = 250 and *t*_de_ = 10^-2^ s. The formation-disruption frequency of the MWEN, *f* is *f* = *N*/*T*_1_ ~ 100 Hz.

These calculations indicate that, under ideal conditions, 250 numbers will be added to the network in every 10^-2^ s interval. When a collapse happens, the entire network will be suddenly broken. The breaking results in all entangled particles within the network simultaneously becoming real ions (Fig. 2). That is, at least one hundred OH^-^ ions and equal H_3_O^+^ ions suddenly appear.

**Fig. 2.**
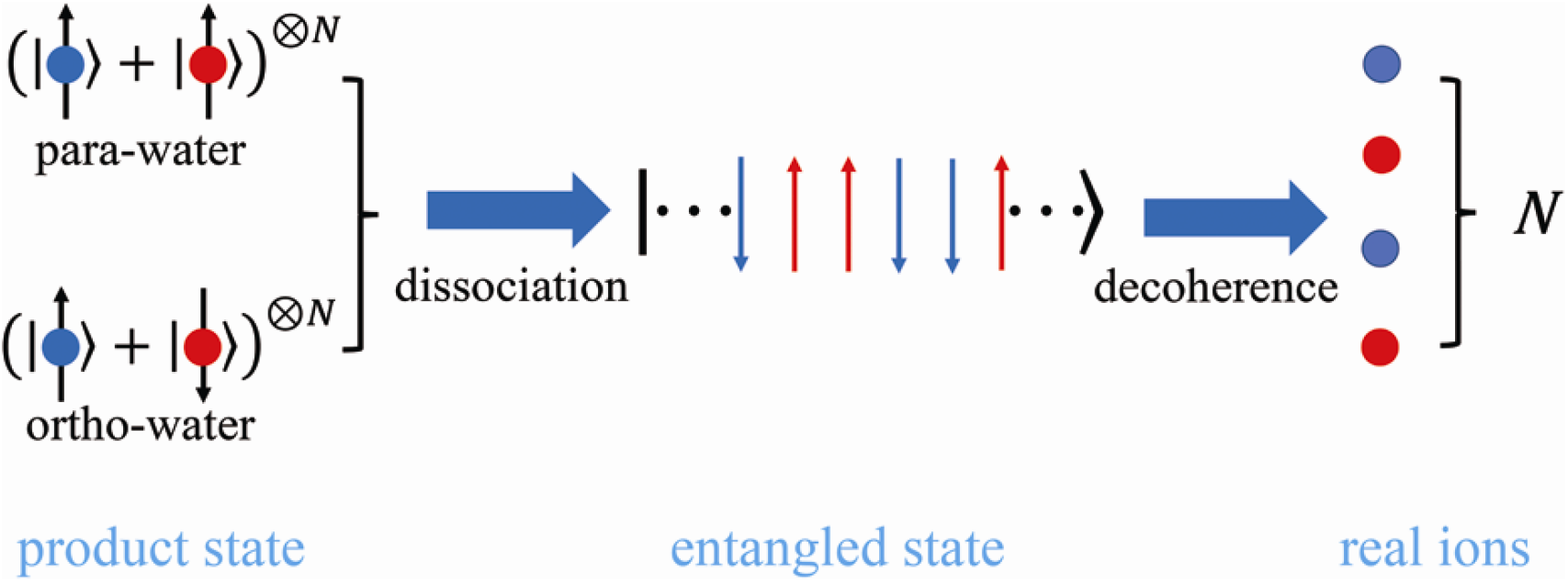
Schematic illustration of Ising model to describe the entanglemnts of the hydronium and hydroxide ions. A symmetric state (*left*) is a product state of the nuclear spins of hydrogen nuclei (within water molecules) which spin-up and -down. An entangled state of hydronium and hydroxide ions (*middle*) is shown where the spins are within a 1-d Ising chain. A collapsed state (*right*) is a state of the real hydronium and hydroxide ions. Red and blue denote different hydrogen nuclei in water molecucles (*left*) and represent different ions (*middle* and *right*). Arrows denote spin directions, and • the ions.

Note that the above calculations considered only a one-by-one pattern of water-dissociation events contributing to the network. Most likely, many water-dissociation events may co-contribute to the network at the same time. An MWEN with co-contributions from many events will be far larger than that from a one-by-one contribution.

Notably, only the water molecules within the hypercoordinations of OH^-^ ions are likely to dissociate (*28*). Therefore, the preexisting OH^-^ ions are the only contributors to the number of an MWEN. Since the disruption of an MWEN will produce at least a hundred-fold more OH^-^ ions than preexisting OH^-^ ions, an MWEN has a great ability to amplify preexisting OH^-^ ions. Increased preexisting OH^-^ ions will promote the formation of a larger MWEN through increased co-contributions. Disruption of a larger MWEN will yield more OH^-^ ions. On the other hand, increased H_3_O^+^ ions will attenuate the amplification by annihilating the preexisting OH^-^ ions.

### MWEN-amplified OH^-^ ions and neuronal excitability

We next linked MWEN-amplified OH^-^ ions to increased neural excitability. For increased neural excitability means more anesthetic requirements to anesthetize humans or animals and *vice versa* (*3*), MWEN amplification capacity may account for the above LORR ED_50_ results.

Of the parameters of an MWEN, the formation-disruption frequency, *f*, is important because *f* may tune the frequencies of conformational fluctuations of proteins and therefore neural excitability. Proteins feature spontaneously intrinsic fluctuations-transitions from one conformational state to another and back, which correspondingly alter their sensitivities to effectors (*29*). OH^-^ ions are hydrophobic (*30-32*), which ensures that OH^-^ ions penetrate the hydrophobic pockets within proteins. Penetration of a swarm of OH^-^ ions, resulting from the disruption of an MWEN, may disrupt the van der Waals forces that normally shape proteins, not only by the mere occupancy but also by the electrostatic forces that the ions carry. An MWEN may thus tune the frequencies of protein conformational fluctuations and therefore their activities or functions. In neurons, the tunning may make ion channels operate cooperatively (*33*). The cooperativity may account for the opposing pH-correlated activities of *N*-methyl-D-aspartate (*34*) (excitatory) and γ-aminobutyric acid (*35*) (inhibitory) receptors, and therefore differing neuronal excitabilities. Higher frequency means that the neuron possesses increased capacity to process information input in the form of action potential frequencies. When a neuron has a low frequency, it will miss some information because some action potentials will be input during the refractory periods of the neuron, that is, the neuron has decreased excitability. Neuronal spiking rates are indicators of neuronal excitability. We next recorded the spiking rates of cultured mouse primary cortical neurons (fig. S2) exposed to the *in vivo* dosages of weak acids and bases used to determine LORR ED_50_ above. We found, as predicted, that weak acids (Fig. 3, A and B) decreased whereas weak bases (Fig. 3, C and D) increased spiking rates, indicating that acids lead to a decrease whereas bases an increase in neuronal excitability.

**Fig. 3.**
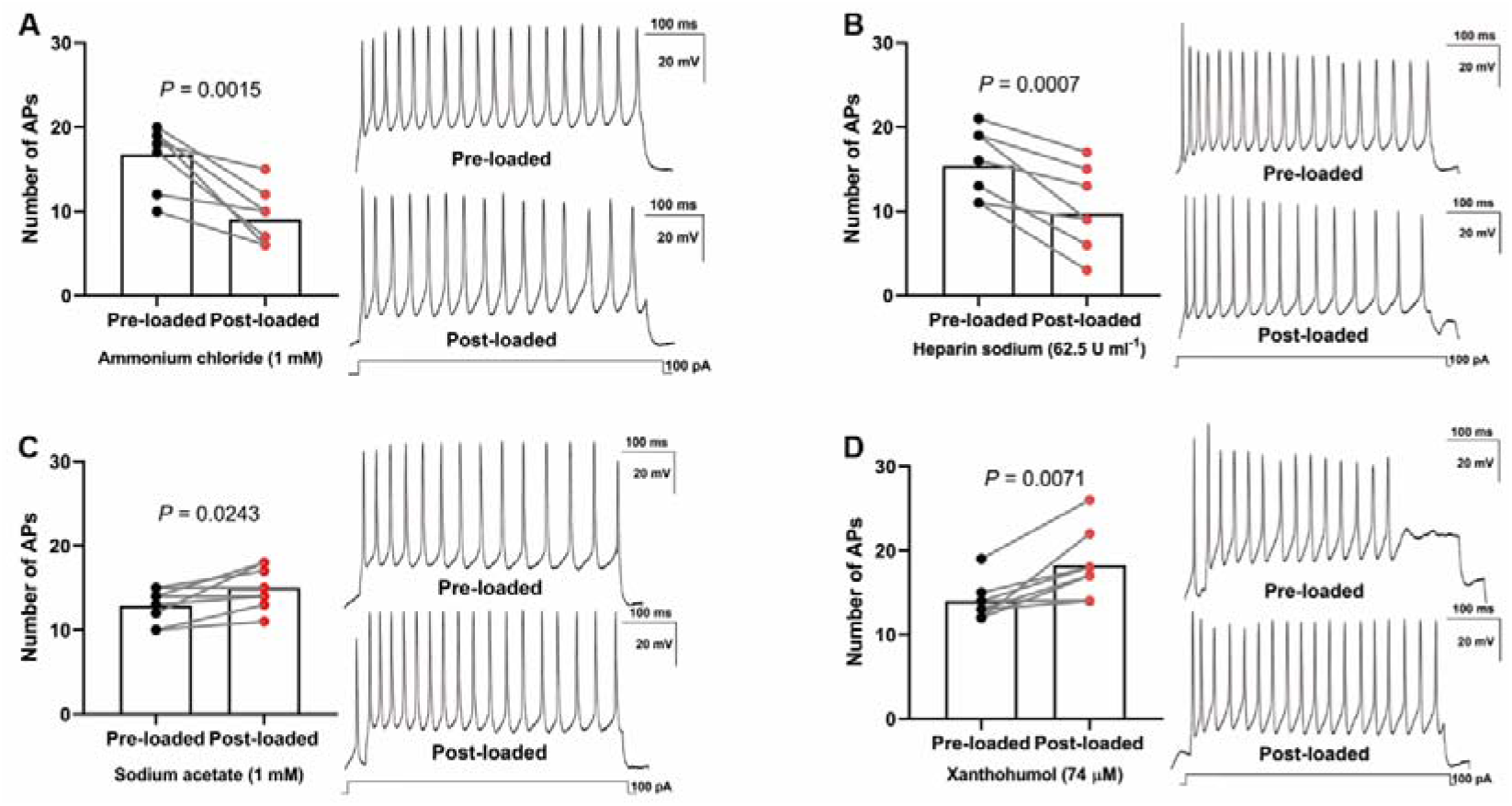
Effects of weak acids and bases on the neuronal spiking rate. Weak acids decreased the spiking rate: **A**, ammonium chloride (pre-SD, 3.69, and post-SD, 3.34; and 95% CI, 13.66 to 19.84 versus 6.209 to 11.79) and **B**, heparin sodium (pre-SD, 3.93, post-SD, 4.86; and 95% CI, 12.09 to 18.66 versus 5.685 to 13.82). Weak bases increased the spiking rate: **C**, sodium acetate (pre-SD, 2.03, post-SD, 2.51; and 95% CI, 12.09 to 18.66 versus 5.685 to 13.82) and **D**, xanthohumol (pre-SD, 2.27, post-SD, 4.03; and 95% CI, 12.10 to 15.90 versus 14.88 to 21.62). The insets (*right*) are the representative recordings of the action potential. In each group, a two-tailed paired *t*-test was used to compare the spiking rate. *n* = 8 cells. AP denotes action potential. The spiking rate is the number of action potentials in 500 ms.

### IAs targeting at formation of an MWEN

Because *f* accounts for neuronal excitability, we next explored whether IAs produce general anesthesia by reducing *f*. Because *f* = *N/T*_1_, to reduce *f*, IAs may ether reduce *N*, prolong *T*_1_ or both. Because IAs such as xenon (*36*) and isoflurane (*37*) at clinically relevant concentrations have negligible effects on *T*_1_, reducing *N* may be the determinant for IAs to reduce *f*.

We used molecular dynamics simulations to calculate the binding energy released when one IA molecule binds with one water molecule (*38, 39*). The binding energy, usually negative (exothermic), represents the water dissociation capacity, which determines *N*. A more negative value means that more energy is needed for the combined water molecule to dissociate and, therefore, that *N* is smaller. Inert (argon, krypton, and xenon) and conventional (isoflurane, desflurane, sevoflurane, and nitrous oxide) anesthetics and a trial anesthetic, F3 (1-chloro-1,2,2-trifluorocyclobutane), were selected (supplementary text), and their binding energies were calculated. We found that the anesthetic potencies of the selected anesthetics correlated remarkably well with their absolute binding energies (Fig. 4, A and B, and figs. S3 and S4 and Table S1), indicating that a reduction in the water dissociation capacity and a consequent weakening of the MWEN formation is responsible for the anesthetic effect of IAs.

**Fig. 4.**
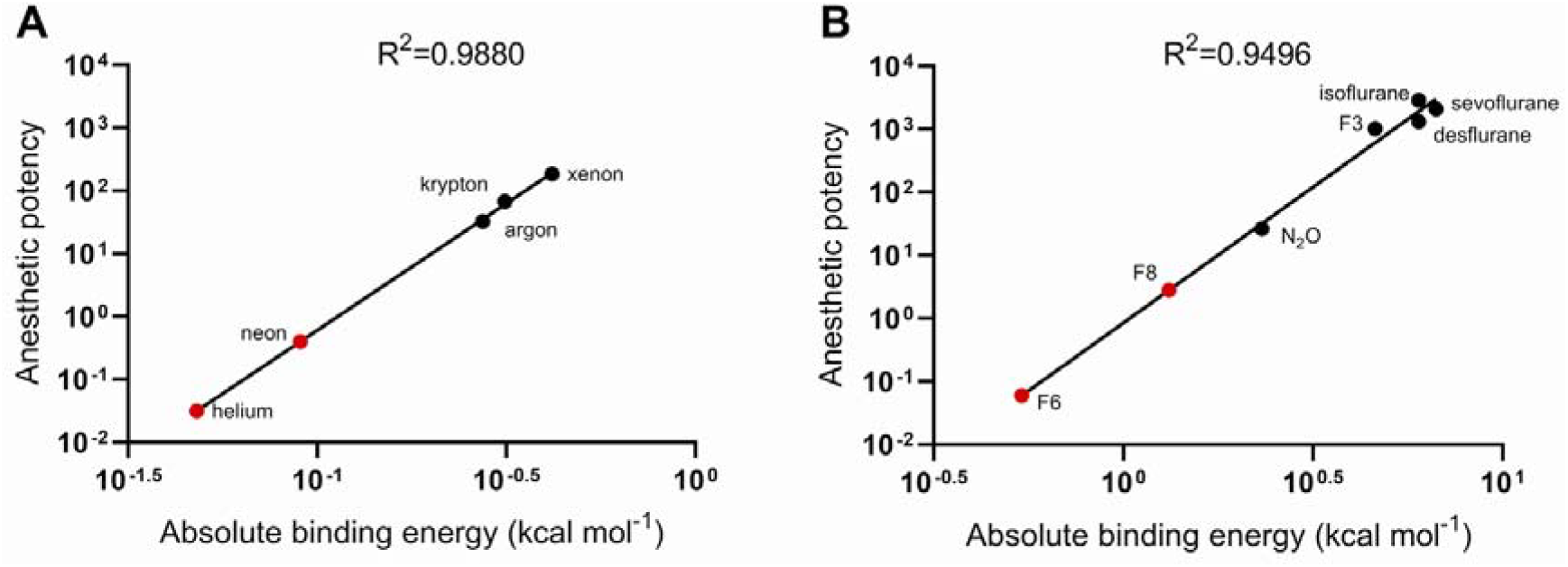
Relationship of the binding energies of the inhaled anesthetics with their anesthetic potencies. The data are plotted as potencies, defined as reciprocals of the aqueous 50% effective concentrations (M) in rats (supplement text). Because the results of different basis sets cannot be compared, the data are presented in two panels. **A**, Inert gases. **B**, Conventional anesthetics and F3. The logarithms of the aqueous 50% effective concentrations for the two classes of inhaled anesthetics correlate well with the logarithms of the absolute values of their binding energies, showing as large values of R^2^. The binding energies of helium and neon (red dots) are plotted in panel **A,** and F6 and F8 (red dots) in panel **B,** to determine their anesthetic potencies by linear regression.

We next used our model to explain some anesthetic phenomena that have not yet been clearly explained. The first is the isotopic dependence of xenon anesthesia (*3*). Because our calculations showed that xenon isotopes have identical binding energies (fig. S3), different effects of xenon isotopes on *T*_1_ may account for the isotopic dependence of xenon anesthesia. Shortening *T*_1_ is possible because ^129^xenon and ^131^xenon have nonzero nuclear spins, which may cross-relax with hydrogen nuclear spins (*40*). We measured the values of 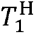 in ^129^xenon-, ^131^xenon- and ^134^xenon-water solutions by using nuclear magnetic resonance imaging and accordingly calculated the *f* ratios. We found that the *f* ratios of ^129^xenon and ^131^xenon to ^134^xenon were quantitively in accord with their LORR ED_50_ values (Table 1 and fig. S5), suggesting that cross-relaxation is the reason for the lower potency of ^129^xenon and ^131^xenon.

**Table 1.** Comparisons of xenon isotopic potencies with their *f* ratios.

| Isotope | LORR ED<sub>50</sub> (%)<sup>a</sup> | $T_1^H$ (ms)<sup>b</sup> | LORR ED<sub>50</sub><br>Ratio<sup>c</sup> | $f$ Ratio<sup>d</sup> |
| --- | --- | --- | --- | --- |
| <sup>129</sup>xenon | 105 | 2357 (144) | 0.69 | 0.68 |
| <sup>131</sup>xenon | 99 | 2583 (195) | 0.73 | 0.75 |
| <sup>134</sup>xenon | 72 | 3448 (202) | 1 | 1 |

<sup>a</sup>Cited from our previous study (3).
<sup>b</sup>Acquired by nuclear magnetic resonance imaging. Mean (SD), $n = 4$ samples for each isotope. The isotopes in this study were the same batches used in our previous experiment (3) to test the LORR ED<sub>50</sub> in mice. No statistical analysis was performed to compare the differences between isotopes.
<sup>c</sup>Calculated using $\frac{\text{LORR ED}_{50}}{^{134}\text{Xe LORR ED}_{50}}$ .
<sup>d</sup>Calculated using $\frac{T_1^H}{^{134}\text{Xe } T_1^H}$ .

Non-immobilizers, such as F6 (1,2-dichlorohexafluorocyclobutane) and F8 (2,3-sdichlorooctafluorobutane), are inhaled compounds that are predicted to be anesthetics from their lipid solubility but do not cause immobility (*12*). Plotting the binding energies of F6 and F8 in Fig. 4B showed that the minimum alveolar concentrations (MACs) of F6 and F8 in rats are 31,436 atm and 4,210 atm, respectively, which are too high to be anesthetics. Helium and neon are non-anesthetics because their MACs are as high as 81,143 atm and 6,382 atm (Fig. 4A), respectively.

Whether IAs have stereoselectivity is under debate. *S*-(+)-Isoflurane was found to be more potent than *R*-(-)-isoflurane when intravenously injected (*41*), but no difference was found when inhaled (*42*) by rats. We found that *S*-(+)-isoflurane and *R*-(-)-isoflurane have identical binding energies (Table S1), suggesting that isoflurane has no anesthetic stereoselectivity, consistent with the fact that IAs act at nonspecific sites.

## Discussion

The present knowledge suggests that anesthesia results from IAs occupying hydrophobic pockets within target proteins (*7, 9*). The mere presence of IAs in hydrophobic pockets, however, presents insurmountable problems. Protein pockets are usually highly selective for ligand shape and size (*2*). Fitting IAs with highly diverse shapes and sizes into structurally selective pockets is problematic. In fact, few proteins possess pockets large enough to accommodate large anesthetics such as halothane (*43*). Furthermore, some compounds occupy the same pockets but do not exert anesthetic effects or even have contrasting effects (*1*). Apparently, components that bridge IAs and protein pockets should be needed. This study indicates that MWEN-amplified OH^-^ ions are these components because our model can easily explain the isotopic dependence of xenon anesthesia, non-immobilizers, and anesthetic stereoselectivity aforementioned. In addition, our model can explain the pressure reversal and why anesthesia populates living organisms. The reversal of anesthesia by high pressure, termed pressure reversal, has been repeatedly observed in different animal species (*44-46*). High pressure enhances water dissociation (*47, 48*), acting as an increase in the preexisting OH^-^ ions in this study. According to our model, if the pressure is high enough, the water dissociation capacity may increase to a level that can fully compensate for the capacity loss caused by the IA, and eventually, the anesthesia is completely reversed. Our model is established on the basis of a causally complete rule-energy minima. Undoubtedly, this causation is applicable to all IAs and to all living organisms-IAs can anaesthetize all living organisms tested thus far, from *Paramecium* to humans to plant (*6*). Taken together, our model offers a unified mechanism for the anesthetic action of IAs.

Due to the many similarities between general anesthesia and sleep (*1, 6*) (the flip side of consciousness), this study suggests that general anesthesia, sleep and consciousness may share a single and simple common mechanism-the MWEN-based amplification of OH^-^ ions. Evidence has indicated that the persistent excitability of neurons results in gradual neuronal acidification (*49*) and that sleep deprivation causes brain acidification (*50*). It is thus reasonable to believe that sleep restores neuronal alkalization that has been consumed by consciousness processing.

## Acknowledgements

We thank Heshui Wu, the MRI Room at Union Hospital, Wuhan, China, for help with MRI testing; Ping Yin, Department of Epidemiology and Biostatistics and State Key Laboratory of Environment Health, School of Public Health, Tongji Medical College, Huazhong University of Science and Technology, Wuhan, China, for statistical assistance; and Hong Ren, Department of Anesthesiology at Union Hospital, Wuhan, China, for assistance in electroneurophysiological experiments.

## Author contributions

H.Q. designed and performed the neuronal culture, MRI, and electroneurophysiological experiments, with assistance from R.C., D.L. and R.Z. N.L. and Q.H. designed the behavioral experiments that were conducted by D.L., R.Z. and R.C. L.Y. and Y.X. designed the molecular dynamic simulations, which were performed by L.Y., Y.X., H.L., N.C. and Q.Z. S.Z. conceptualized and supervised the study, analyzed the data, and wrote the manuscript with input from all authors.

## Data and materials Availability

All data are available in the main text or the supplementary materials. All materials are available upon request from S.Z.

## Materials and Methods

### Animals

C57BL/6 male mice (7 weeks; weight range, 22-25 *g*) and C57BL/6 female mice (postnatal days, 18-19) were ordered from Beijing Vital River Laboratory Animal Technology Co., Ltd. ([jing]2016-0006; China). The animals were bred in a temperature- and humidity-controlled room with a 12-h light/dark cycle. The animals had free access to standard mouse chow and tap water before the experiments. The animals used for behavioral experiments were housed in the room for a week. All animal operations and experimental protocols conformed to the US National Institutes of Health Guide for the Care and Use of Laboratory Animals (NIH Publications No. 8023, revised 1978) and were approved by the Institutional Animal Care and Use Committee (approval No: S164) at Tongji Medical College, Huazhong University of Science and Technology.

### Agents

Sevoflurane was purchased from Maruishi Pharmaceutical Co., Ltd. (Chouku, Osaka, Japan). Heparin sodium was purchased from Changzhou Qianhong Biochemical Pharmaceutical Co., Ltd. (Changzhou, China). Xanthohumol was obtained from J&K Scientific Ltd. (Beijing, China). ^129^Xenon, ^131^xenon and ^134^xenon gases were purchased from NUKEM Isotopes Imaging GmbH (Alzenau, Germany) and all had a purity ≥ 99.99% and an abundance of 93.5%.

Culture media, including Dulbecco’s modified Eagle’s medium/F12 (1:1) (DMEM/F12), fetal bovine serum (FBS), 0.25% trypsin solution, calcium- and magnesium-free Hank’s balanced salt solution (CMF-HBSS), phosphate-buffered saline (PBS), neurobasal medium and B-27 supplement, were purchased from Life Technologies Gibco^®^ BRL (Grand Island, NY, USA). Poly-D-lysine, DNase I, penicillin/streptomycin, and L-glutamine were purchased from Sigma-Aldrich Co., Ltd. (St. Louis, MO, USA). All other reagents of analytical grade were purchased from Sinopharm Chemical Reagent Co., Ltd. (Shanghai, China).

### Solution preparations

All solutions were prepared immediately prior to the experiments. For the animal behavioral experiments, 1 mM sodium acetate (CH_3_COONa), 1 mM ammonium acetate (CH_3_COONH_4_), 1 mM ammonium chloride (NH_4_Cl), heparin sodium 62.5 U/ml and 74 μM xanthohumol were prepared by dissolving in normal saline. All solutions were adjusted to an osmolality of 300 ± 10 mOsm with NaCl or tri-distilled water. The solutions were filtered through a 0.22-μm filter (Millipore, Bedford, MA, USA) prior to use.

### LORR ED_50_ determination

To determine the LORR ED_50_, sixty C57BL/6 male mice (8 weeks old) were divided randomly into saline, ammonium chloride, sodium acetate, ammonium acetate, heparin sodium, and xanthohumol groups, with 10 mice in each group. The LORR ED_50_ of sevoflurane in mice was determined according to our previously described method (*10*) with slight modifications. Briefly, the basal and drug-loaded LORR ED_50_ of each mouse was determined, and all the values were compared. For the basal LORR ED_50_ determination, mice were individually placed in an isolated plastic mesh V-shaped trough fixed in a clear plastic chamber (205×134×69 mm) with an electrical fan to mix gases. One side of the chamber was connected to a sevoflurane vaporizer (Aika, Ichikawa Shiseido, Tokyo, Japan). The other side was connected to an infrared gas monitor (Bene View T5, Mindray Bio-Medical Electronics, Shenzhen, China) to measure the sevoflurane, oxygen and carbon dioxide concentrations in real time. The monitor can measure the sevoflurane concentration with a precision of 0.01%.

When a mouse was placed in the chamber, pure oxygen was immediately supplied at a rate of 600 ml/min. When the chamber’s oxygen concentration increased to 99%, sevoflurane gas mixed in pure oxygen was provided by the vaporizer. The initial sevoflurane concentration in the chamber was 1.00%, which was maintained for 15 minutes to equilibrate the mouse with sevoflurane gas. Then, the chamber was rotated 180° to place the mouse on its back in the V-trough, and its righting reflex was observed. LORR was defined as the supine mouse unable to turn itself onto all four paws three times within 1 min. According to the mouse’s righting reflex, a stepwise increase or decrease of 0.10% sevoflurane in the chamber was applied. Specifically, if the mouse’s righting reflex disappeared, the sevoflurane concentration was decreased 0.10%; otherwise, it was increased 0.10%. After 15 minutes of equilibration at each sevoflurane concentration, the mouse’s righting reflex was observed again. The LORR ED_50_ was the average of the two critical sevoflurane concentrations at which the mouse either lost or regained its righting reflex (*10*). All righting reflexes were observed by a trained observer who was unaware of group allocation. All determinations were made between 08:00 and 18:00.

After basal LORR ED_50_ determination, the mouse underwent intracerebroventricular cannulation following our previously described method (*10*). After 1 day of recovery, the LORR ED_50_ of the mouse was determined again. Briefly, the mouse was sedated with 1% sevoflurane. Then, saline, ammonium acetate (1 mM), sodium acetate (1 mM), ammonium chloride (1 mM), heparin sodium (62.5 U/ml), or xanthohumol (74 μM) was injected into the lateral cerebral ventricle by a syringe pump system (RWD Life Science Co., Ltd., Shenzhen, China). All agents were warmed to 37 °C before infusion. The intracerebroventricular injection was performed automatically by the pump at a rate of 1 μl/min. A total volume of 2 μl was injected. After injection, sevoflurane was discontinued, and the mouse could recover. When the mouse could move freely, the LORR ED_50_ of sevoflurane in the animal was determined.

### Cortical neuron culture

Primary culture of mouse cortical neurons was performed using a described method (*51*) with some modifications. In brief, under sevoflurane anesthesia, mouse embryos were removed from timed pregnant mice at embryonic day 18. The embryos were killed by decapitation quickly in ice-cold CMF-HBSS solution. Brains were rapidly dissected, and the cortex was cleared from the meninges and isolated under a dissection microscope. Cortices were collected in ice-cold CMF-HBSS with 0.2% D-glucose and 1% penicillin-streptomycin solution and were then homogenized. Next, the tissues were digested with 0.125% trypsin and 100 μg/ml DNase I in CMF-HBSS for 15 min at 37 □. After filtering the tissues through a sterile nylon sieve with a pore size of 70 μm, cells were collected by centrifugation for 5 min at 80 *g*. The cells were suspended in DMEM/F12 containing 10% FBS and 1% penicillin-streptomycin at 5×10^5^ cells per ml and subsequently inoculated at 400 μl per well into 24-well culture plates containing coverslips (Flexcell International Co., Hillsborough, NC, USA). The coverslips were coated with poly-D-lysine (100 μg/ml). After 4 h of incubation in a humidified chamber provided with 5% CO_2_ and 95% air at 37 □, the cells were washed three times with PBS and replaced with 400 μl neurobasal/B27 medium supplemented with 500 μM L-glutamine. Three days later, the culture medium was replaced with fresh culture medium lacking L-glutamine. Half of the medium was then replaced every 3 days. The cells were used for experiments after 11-14 days of culture.

### Electrophysiological recording

Cultured primary cortical neurons were used to perform electrophysiological recording (*52*). Neuronal action potentials were obtained by using current-clamp mode. Cortical neurons cultured on coverslips were placed in the prefusion chamber with an extracellular solution with the following composition: 140 mM NaCl, 5 mM KCl, 1 mM MgCl_2_, 2 mM CaCl_2_, and 10 mM HEPES (pH adjusted to 7.3-7.4 with NaOH). The pipette solution consisted of 130 mM KCl, 10 mM NaCl, 1 mM MgCl_2_, 10 mM EGTA, 10 mM HEPES, 10 mM glucose, and 2 mM ATP-Na_2_ (pH adjusted to 7.2 −7.3 with KOH). Patch pipettes were pulled to a tip resistance of 6-8 MΩ using borosilicate glass capillary tubes with a P-1000 micropipette puller (Sutter Instrument, Novato, CA, USA). Current clamp recording was performed using a Multiclamp 700B amplifier (Axon Instruments, Foster City, CA, USA) equipped with a Digidata 1440 data acquisition system and pCLAMP 10 software (Axon Instruments, Foster City, CA). Data were digitized at 10 kHz. The cortical neurons were held at 0 pA. The action potentials were elicited using a depolarizing current of 100 pA (500 ms) to obtain the basal value. The cells were then exposed to dosages of the weak acids and bases (used to determine the LORR ED_50_ in the mice). The tested compounds were added to the extracellular solution and delivered through a perfusion system (Minipuls 3, Gilson International France SAS, Villiers-le-bel, France) at 37 °C and a rate of 3 ml/ min for 2 min. Then, the neurons were perfused with extracellular solution (37 °C) without the tested compounds for 2 min to imitate drainage of the drug injected into cerebrospinal fluid. Drug-induced changes in the number of action potentials for the cells were elicited and recorded at 2 min of removal. All solutions were warmed and kept at 37 °C.

### 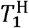 acquisition

Oxygen was removed from the samples of xenon-water solutions by the following procedure. A 250-ml clear glass bottle was filled with 200 ml tri-distilled water and then covered. A 21G needle was inserted into the cover of the bottle. The needle was kept open to let air out. The bottle was then warmed in a water bath at 60 □ overnight. Three bottles were prepared. After being warmed up, the bottles were first separately pressurized with ^129^xenon, ^131^xenon, and ^134^xenon gases for one minute, and then the gases were quickly flushed out. The pressurization-flushing procedure was repeated three times before the bottles were finally sealed. Water with an oxygen partial pressure as low as undetectable by an ABL800 FLEX blood gas analyzer (Radiometer Medical ApS, Brønshøj, Denmark) was sampled. The bottles were then separately filled with ^129^xenon, ^131^xenon, and ^134^xenon to 1 bar and sealed to let the xenon mix with the water overnight. Then, 30 ml xenon-water solution was drawn into a 50-ml plastic syringe and sealed for the 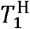 test. Four samples were prepared for each isotope.

Magnetic resonance imaging (MRI) was performed to measure 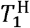 values at the MRI Room of Department of Radiology at the Union Hospital of Tongji Medical College of Huazhong University of Science and Technology, Wuhan, China. We used a super-high-field superconducting 3.0 Tesla MRI scanner (Magnetom Trio, Siemens Healthcare Solutions, Inc., Erlangen, Germany) to obtain thin-section planar scans (1 mm section thickness) with a 32-channel head coil. All samples were measured at the same time. The imaging protocol consisted of axial T1-weighted imaging (T1WI) of fast spin echo sequences. T1WI as 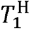 was acquired with the following parameters: repetition time, 350 ms; echo time, 11 ms; field of view, 4.0 × 4.0 mm; matrix size, 256 × 256; and slice thickness, 1 mm.

### Molecular dynamics simulations

For inert gases, the geometries of XH_2_O (X = helium, neon, argon, krypton, ^129^xenon, ^131^xenon, or ^134^xenon) complexes were optimized, and their binding energies were calculated using the QZVPP basis set at coupled-cluster singles and doubles with perturbative triple correction [CCSD(T)] at the def2-QZVPP level of theory (*53*) using the ORCA (*54*) suite of programs (version 4.2.0).

For conventional anesthetics, F3 and non-immobilizers, the geometries of AH_2_O (A = nitrous oxide, isoflurane, sevoflurane, desflurane, F3, F6, or F8) complexes were fully optimized, and their binding energies and harmonic frequencies were calculated at density functional theory (DFT) (*55*) and the B3LYP/def2-TZVPPD level of theory, which were implemented in Gaussian 16W (version C.01 for Windows, Gaussian, Inc., Wallingford, CT, USA) suite of programs. For each complex, the harmonic vibrational frequencies and binding energies of all configurations were calculated. If configurations had imaginary frequencies, they were excluded from subsequent data analysis. For each complex, the configuration with the lowest binding energy was selected for subsequent data analysis according to the rule of energy minima. Avogadro (version 1.2.0) software (*56*) for Windows was used for molecular modelling and mapping.

### Statistics

Sample sizes were predetermined, and randomization and blinding were performed according to our previous study (*10*). Data are presented as the means ± standard deviations. Single comparisons of within-group measures of the LORR ED_50_ and the number of action potentials were made by a two-tailed paired Student’s *t*-test after the data were examined for normality of distributions using the Shapiro-Wilk test. The binding energy and 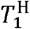 data were analyzed with linear regression. All analyses were performed using SAS version 9.4 (SAS Institute, Cary, NC, USA).

### Supplementary Text

#### Drug-induced pH changes in the mouse brain

Take sodium acetate as an example. We found that the pH of 1 mM sodium acetate at room temperature (25 □) was 7.75, as measured with a pH meter. The pH of cerebrospinal fluid, pH_c_, is 7.35, and the total mouse brain volume (cerebrospinal fluid volume with cell component volume) is 462 μl (*13*). The injection of 2 μl of sodium acetate will replace 2 μl of cerebrospinal fluid. The H^+^ concentration of the mouse brain after injection of sodium acetate, [H^+^]_ac_, is [H^+^]_ac_ = [2×10^-7.75^ + 460×10^-7.35^]/462 = 10^-7.351^ M. pH_ac_ = -log[H^+^]_ac_ = 7.351. _Δ_pH = pH_ac_ - pH_c_ = 0.001. That is, a 2 μl injection of 1 mM sodium acetate into mouse cerebrospinal fluid caused a net change in the pH of the mouse brain of approximately 0.001. The other three drugs would result in net pH changes similar to that from sodium acetate.

Because we are unable to know the average pH value in the brain of the intact mice, we take cerebrospinal fluid pH as brain pH into the above calculation. The intracellular pH is lower than that of cerebrospinal fluid. One can see that _Δ_pH is also approximately 0.001 if pH value of 7.00 is taken into calculation. If the buffer systems of the brain and physiological drainage of cerebrospinal fluid are taken into consideration, the _Δ_pH may be even less. A pyramidal cell, the largest cell of hippocampal neurons in the CA1 layer, is 1,660 μm^3^ in cell volume (*57*). Inputting an intraneuronal pH of 7.00 and the fact that 60% of the cell volume is water into the calculation (*58*), we obtained 6,000 free H_3_O^+^ ions in the cytoplasm of the cell. A decrease of 0.001 pH units in the cytoplasm of the cell after intracerebroventricular injection of ammonium chloride, for example, is equivalent to a net increase of 137 free H_3_O^+^ ions in a pyramidal cell.

#### Inhaled anesthetics selected

Inhaled anesthetics may have mediatory and modulatory effects. Mediatory effects are the mechanism-involved (direct) effects of inhaled anesthetics. Modulatory effects may modulate the aesthetic potency of inhaled anesthetics through non-mechanistic (indirect) routes (*59*) such as directly binding with lipids, receptors, or ion channels. For a given inhaled aesthetic, its potency, such as its MAC, is the sum of its mediatory and modulatory effects. The modulatory effects may thus strongly affect mechanistic studies because mechanistic studies rely on only mechanism-involved MACs. This study is a mechanistic study. We thus had to select anesthetics with known minimal modulatory effects to plot Fig. 4 in the main text.

Different animal species may have different MACs. For a given animal species, different laboratories may obtain different MACs derived from different experimental paradigms involving different nociceptive stimuli. As such, in this study, the MACs of the inhaled anesthetics were cited from one laboratory (Department of Anesthesia and Perioperative Care, University of California, San Francisco, CA, USA) in one animal species, Sprague-Dawley rats. If discrepancies appeared, the most recently published data were cited. Finally, the MACs in Sprague-Dawley rats for inert gases (argon, krypton, and xenon) (*60*), nitrous oxide (*61*), conventional anesthetics (isoflurane, desflurane, and sevoflurane) (*62*), and F3 (1-chloro-1,2,2-trifluorocyclobutane) (*12*) were cited. The MACs were converted into aqueous ED_50_ values at 37 □ using Henry’s law (*63*) to plot Fig. 4 in the main text.

**Fig. S1.**
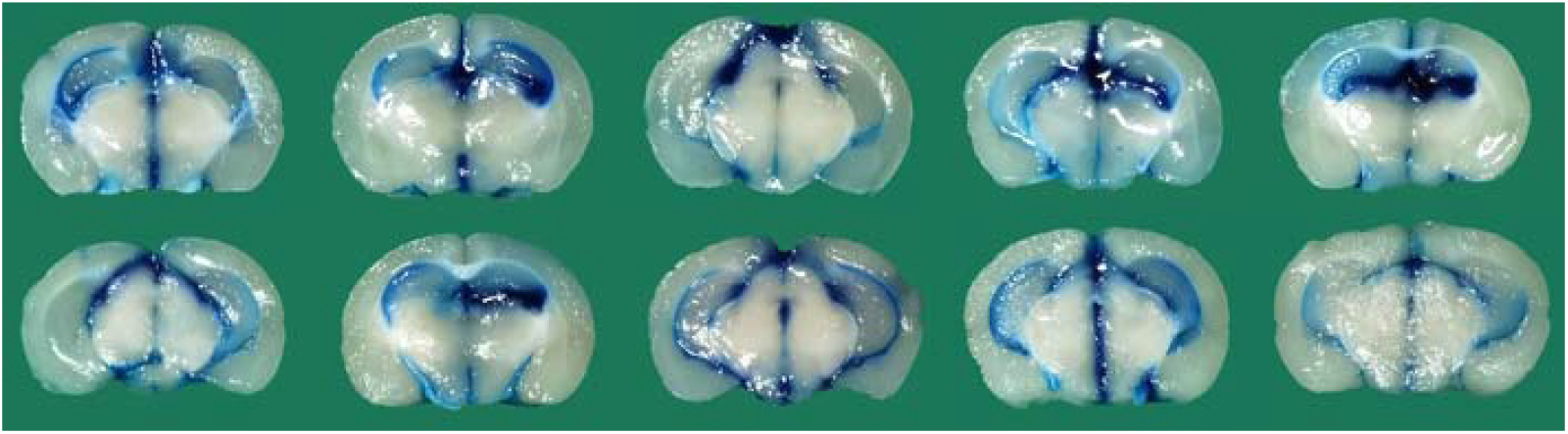
Diffusion of Evans blue throughout mouse ventricular system. After righting reflex testing, 10 mice in the saline group were anesthetized with sevoflurane and then received injection of Evans blue 2 μl into lateral ventricle through ventricular cannula, as the injection procedure as that of tested drugs. The mice were then decapitated, and the brain was removed to check the diffusion of Evans blue (*64*). The photos show that Evans blue diffuses well throughout all mouse ventricular systems, indicating that drug-induced anesthetic effects in Fig. 1 were resulted from global rather than local brain actions of the drugs. After righting reflex testing, all 50 mice in the other five groups were weighed and then housed for a week. The mice were then weighed again. For each mouse, its post-housing weight was compared with its pre-housing weight. If there was a weight loss, the mouse may suffer possible neural pathologic damage (*65*). We did not find any mouse with weight loss.

**Fig. S2.**
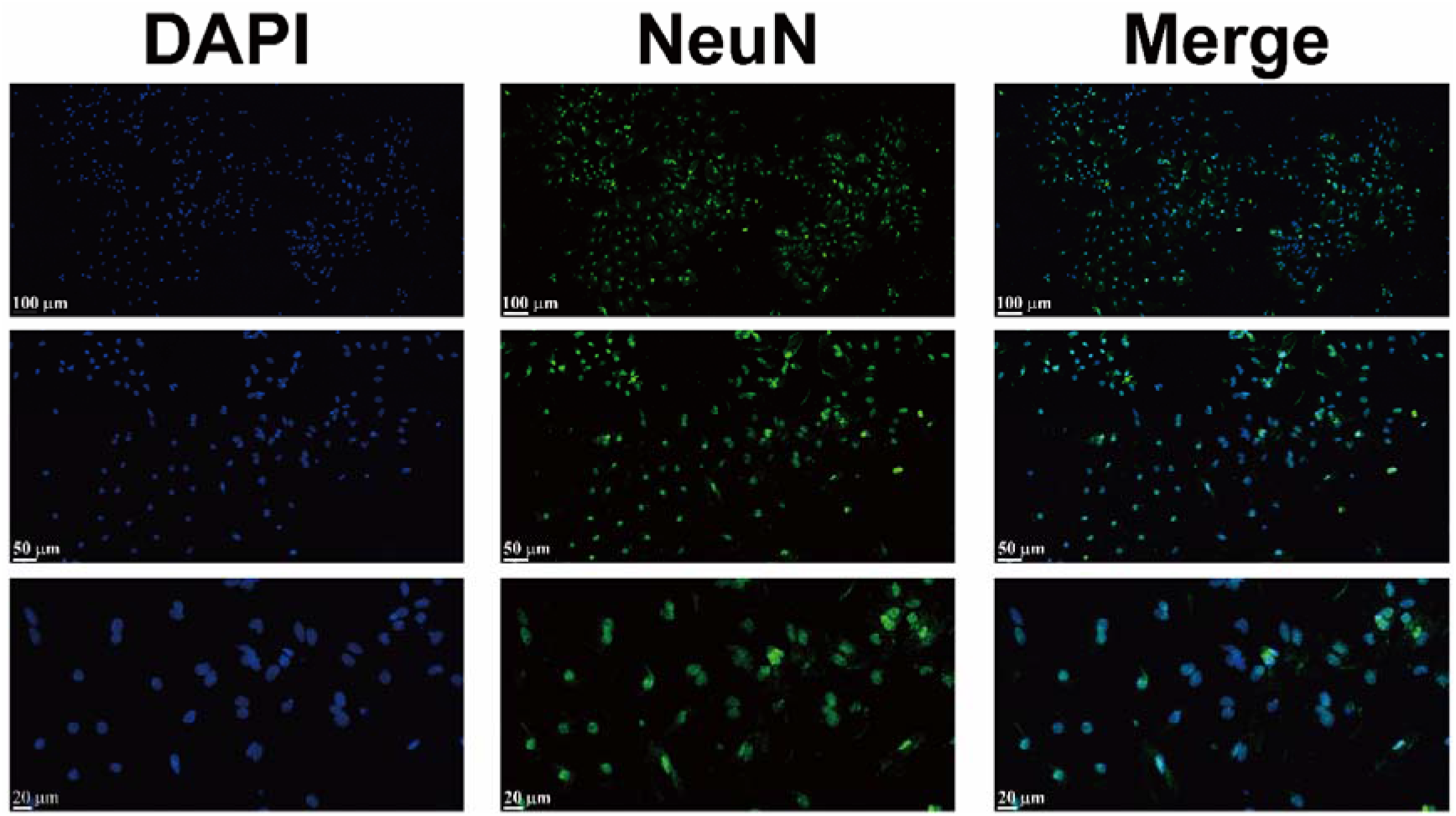
Immunofluorescence images for visualization of primary neurons at 14 days in vitro. Representative images of C57BL/6 primary cortical neurons cultured for 14 days. The cell nuclei were stained with primary anti-NeuN antibody and DyLight 488-labelled secondary antibody (green) and then counterstained with DAPI (blue). The images were then merged to visualize neurons (double-stained). Three scaled (scale bars, 20 μm, 50 μm and 100 μm) images are shown.

**Fig. S3.**
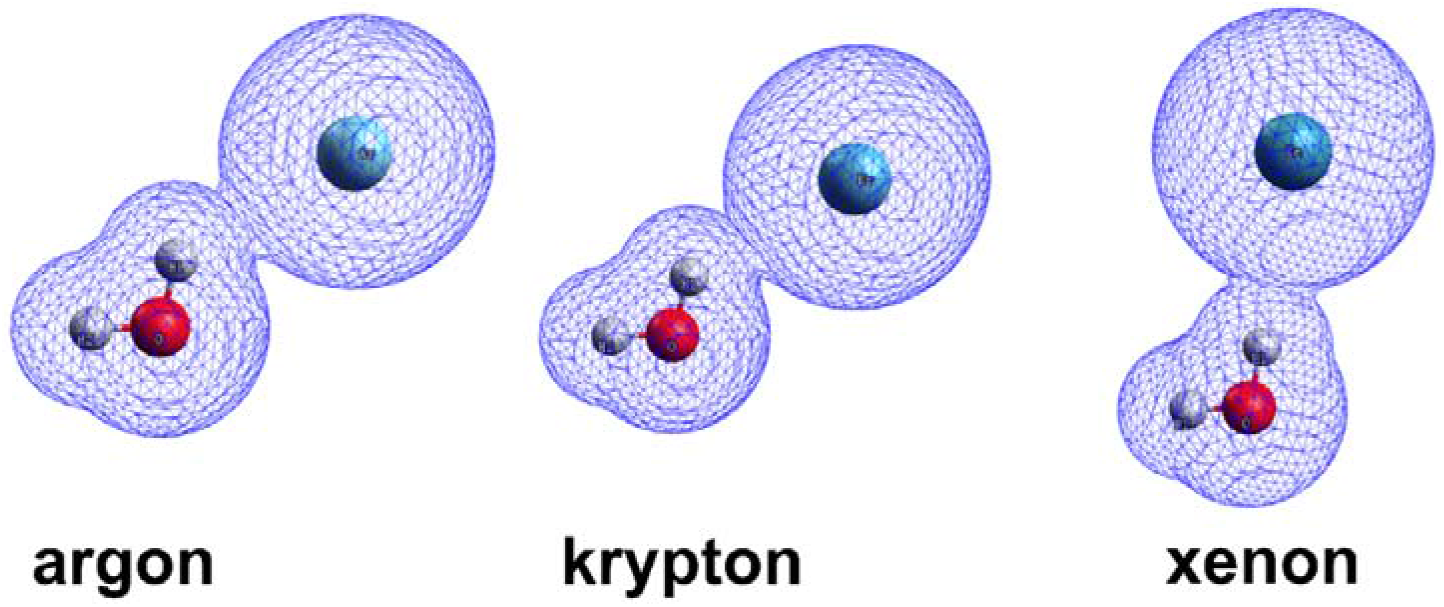
Structures of inert anesthetics with water molecules. The geometries of argon, krypton and xenon combined with one water molecule are shown. The molecular structures are depicted with balls and sticks, and the van der Waals surfaces are shown with thin blue lines around the molecules. The geometries and binding energies of argon, krypton, and xenon were obtained by molecular dynamics simulations using the QZVPP basis set at coupled-cluster singles and doubles with perturbative triple correction [CCSD(T)] at the def2-QZVPP level of theory. The binding energies of helium and neon are −0.048 and −0.090 kcal/mol, respectively, which are used to plot Fig. 4A to obtain their MACs using linear regressions. ^129^Xenon, ^131^xenon, and ^134^xenon have identical binding energies of −0.4177 kcal/mol.

**Fig. S4.**
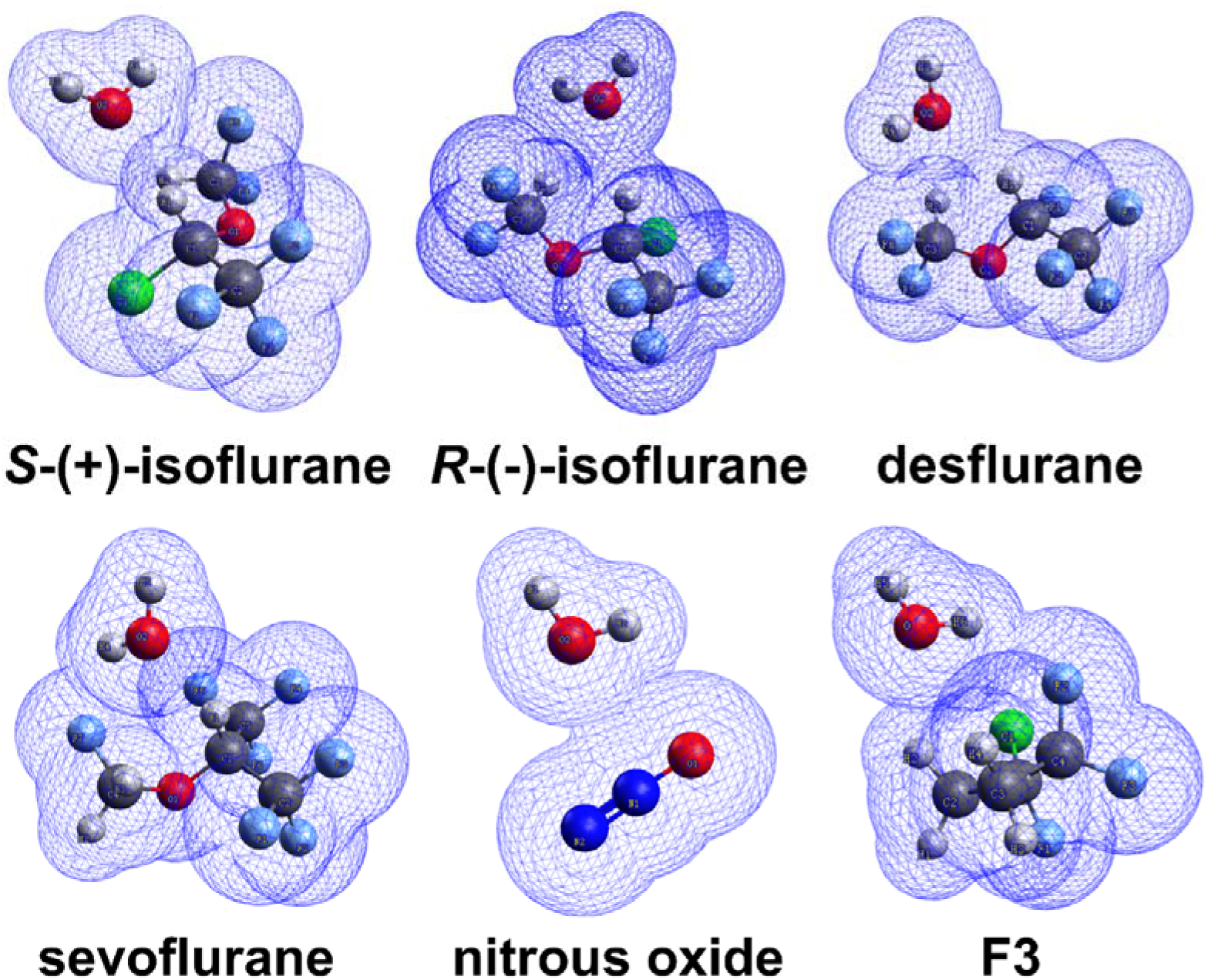
Structures of conventional anesthetics with water molecules. The geometries of isoflurane, desflurane, sevoflurane, nitrous oxide and F3 combining with one water molecule are shown. The molecular structures are depicted with balls and sticks, and the van der Waals surfaces are shown with thin blue lines around the molecules. The geometries and binding energies were obtained by using molecular dynamics simulations at the density functional theory (DFT) and B3LYP/def2-TZVPPD levels of theory. Note that *S*-(+)-isoflurane and *R*-(-)-isoflurane are structurally symmetrical. Because of the symmetry, the two enantiomers have the same binding energy.

**Fig. S5.**
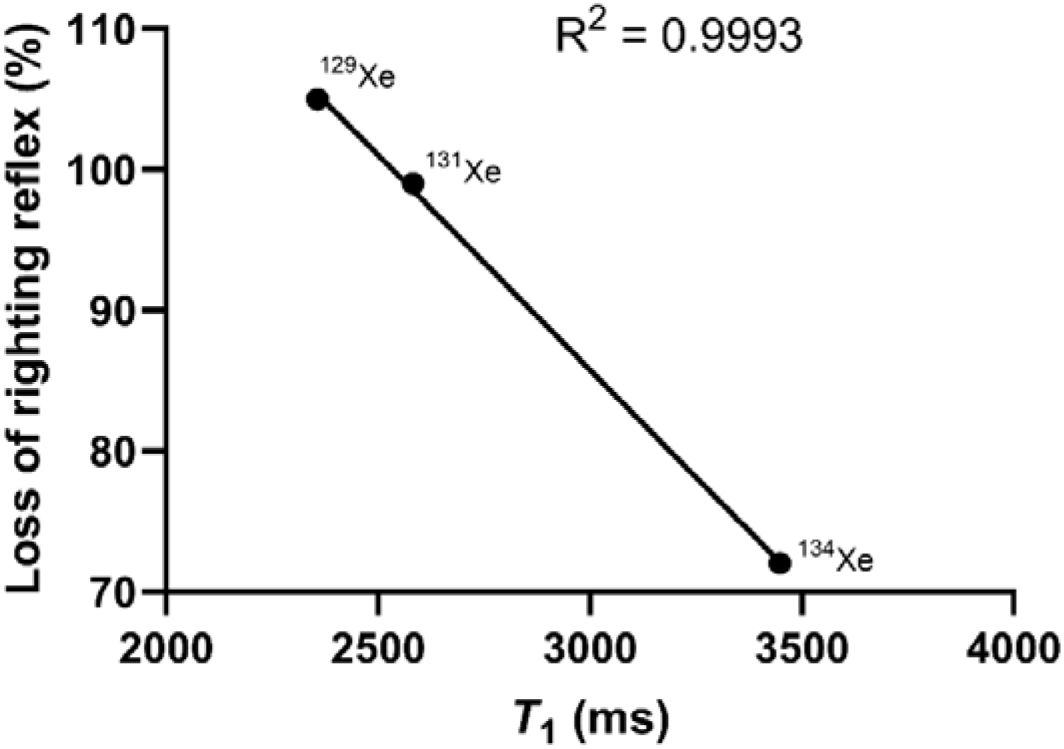
Correlation of the aesthetic potencies of xenon isotopes with *T*_1_. The data in Table 1 in the main text were used to plot this figure.

**Table S1.**
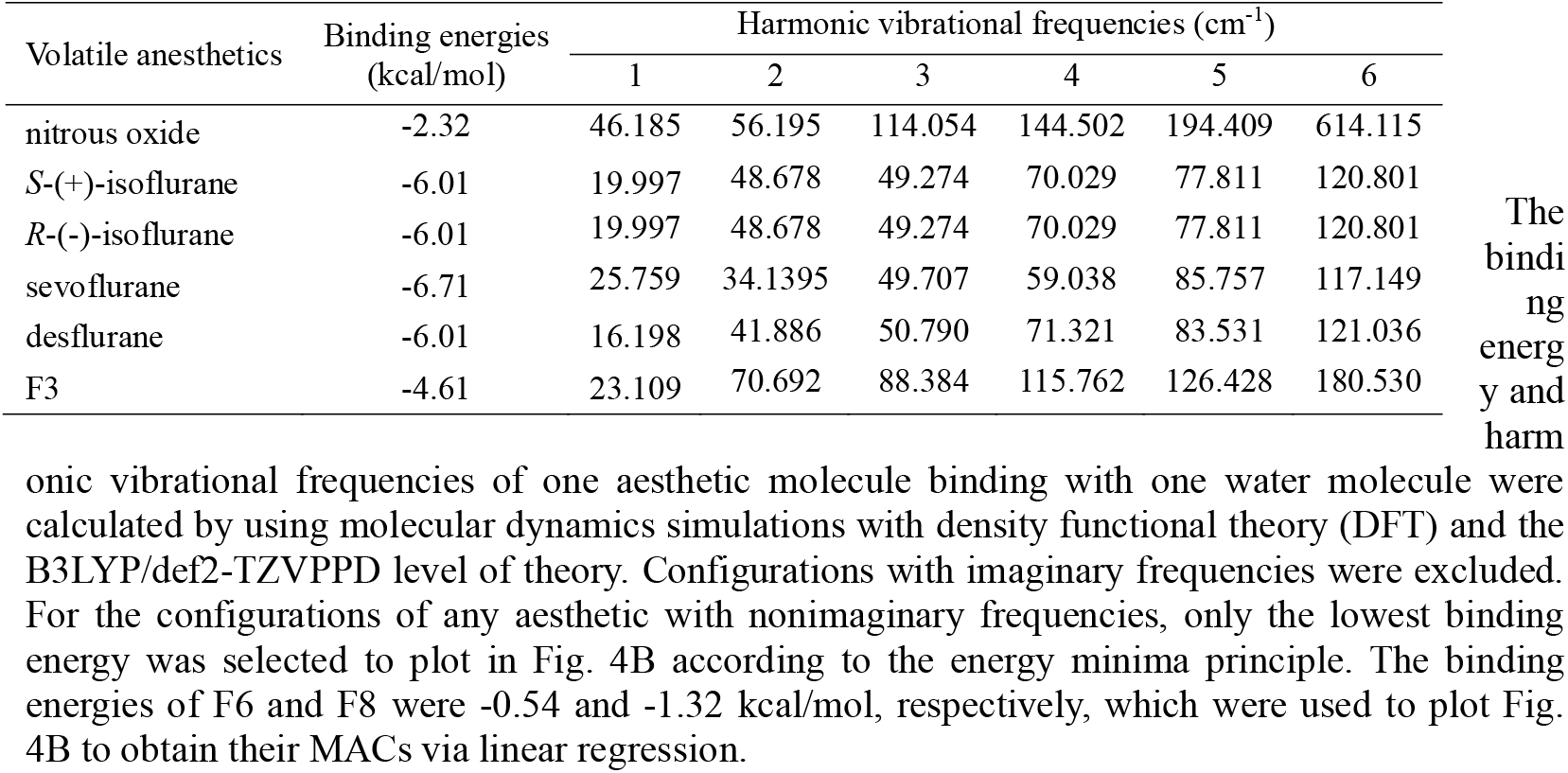
The binding energies and harmonic vibrational frequencies between conventional anesthetics and water molecules.

Different animal species may have different MAC values. For a given animal species, different laboratories may obtain different MAC values derived from different experimental paradigms involving different nociceptive stimuli. As such, in this study the MACs of inhaled anaesthetics were cited from one laboratory (Department of Anaesthesia and Perioperative Care, University of California, San Francisco, CA, USA) in one animal species, Sprague-Dawley rats. If discrepancy appeared, the most recently published data were cited. Finally, the MACs in Sprague-Dawley rats to inert gases (argon, krypton, and xenon)^60^, nitrous oxide^61^, conventional anaesthetics (isoflurane, desflurane, and sevoflurane)^62^, and F3 (1-chloro-1,2,2-trifluorocyclobutane)^9^ were cited. The MACs in partial pressure were converted into aqueous ED_50_ values at 37 □ using Henry’s law^63^ to plot Fig. 3 in the main text.

